# Amino acid clustering and epistasis guide sequence-based prediction of phenotype modulation

**DOI:** 10.64898/2026.11.06.764933

**Authors:** Ayelén S. Hernández Berthet, Ignacio E. Sánchez, Martín E. Noguera, Ernesto A. Roman

## Abstract

Protein sequences encode the molecular information necessary for their biological function. As functional adaptation often emerges from combined effects of multiple positions, understanding how sequence variation gives rise to functional differences remains a central challenge. Here, we developed an improved strategy to identify phenotype modulating positions and use them to predict phenotypes, by relating amino acid identity to experimentally measured quantitative properties across families. Using family-specific sequence-phenotype datasets, our results show that epistasis plays a crucial role in phenotype modulation. Position specific amino acid clustering further improved performance indicating that key functional information is often captured by broad physicochemical amino acid classes, with hydrophobicity playing a crucial role in phenotype modulation. Our identified phenotype modulating positions involve both proximal and distal sites relative to functional sites. While the former is captured by individual positions, the latter is captured when epistasis is taken into account. Comparison with experimentally identified positions showed partial recovery of known functionally relevant residues, while at the same time revealing candidate positions that may contribute to phenotypic variation. Taken together, these results suggest that quantitative phenotypes are often governed by a network of interacting amino acid residues and may be engineered by a limited amount of sequence modifications. This finding highlights the potential of our framework to deepen our understanding of protein function, evolution, and adaptation, providing new targets for the rational modulation and engineering of protein phenotypes.

## Introduction

Proteins encode information such as its three-dimensional architecture, conformational dynamics and their biological function within their amino acid sequences. Whereas structure prediction has been largely solved by the use of deep learning from a practical viewpoint [1,2], prediction of molecular function and phenotypes remains a substantially more complex task. Many phenotypes are measurable quantities whose magnitude varies among homologous proteins and are encoded in their sequences. Thus, understanding how sequence variation affects their magnitude is of fundamental importance as it illuminates the evolution of these phenotypes and should enable quantitative predictions. Prediction of quantitative phenotypes bears direct practical relevance for protein engineering aiming to produce molecules that function under specific conditions of biotechnological or industrial interest.

Quantitative phenotypes which we will collectively refer to as phenotypes, span a broad spectrum. At one end are protein-intrinsic properties, such as ligand-binding affinities, thermostabilities, or catalytic parameters, which arise directly from the protein sequence. At the other end are organismal or contextual phenotypes, such as the optimal growth temperature, that do not depend on a single protein but on an interaction network that leaves adaptive signatures in the involved sequences. Advances in experimental approaches have made it possible to experimentally explore the relation between sequence and quantitative phenotypes. This was studied at scale using deep mutational scanning [3,4], which enables the simultaneous characterization of large numbers of variants within a single experiment. However, these experiments remain resource-intensive and are typically limited to a small number of proteins and variants, hindering the broader exploration of sequence space across homologous proteins and highlighting the need for predictive tools. Most of the existing ones have been developed under conceptually distinct computational frameworks and can be divided between the ones not using preexisting quantitative phenotype information and those that do. The former includes approaches that infer functionally relevant positions from sequence variation including conservation-based [5–7], evolutionary trace [8,9], and statistical coupling analysis [10,11]. The latter include phylogenetic comparative approaches [12,13], like phylogenetic imputation [14], that link sequence changes to phenotype changes in a statistically principled way, without identifying the positions responsible for its modulation. Machine learning approaches currently define the state of the art in predictive accuracy, with examples including predictors of catalytic constants such as DLKcat [15], MPEK [16], and CatPred [17], as well as predictors of other quantities such as the absorption maxima of rhodopsins [18,19] and the optimal growth temperature of organisms [20]. While achieving high predictive accuracy these approaches typically require large training datasets, offering limited mechanistic insight into the sequence determinants, and providing limited interpretable guidance for rational protein engineering. In contrast to these property-specific predictors, one strategy toward a general framework applicable to diverse properties and protein families is transfer learning, where, as an example, a ridge regression method is applied to protein sequence embeddings obtained from a pre-trained protein language model without the need to design a dedicated architecture for each new property [21]). Straightforward, interpretable sequence-phenotype correlation methods [22–25] link positional sequence divergence with differences in quantitative phenotype values or specificity across protein sequence pairs, allowing the identification of positions whose residue identity co-varies with the magnitude of the property of interest. These interpretable methods based on positional sequence analysis face two main challenges. First, sequences with documented phenotype values for a protein family are required. Second, a sensitive distance metric must be applied to capture the position divergence, with the additional challenge of accounting for non-linear relationships between positions. Together, these gaps underscore the need for approaches that are simultaneously interpretable, mechanistically informative, and applicable in data-limited regimes.

Here, building on our previous work, we present an improved protocol to identify phenotype-modulating sites and use them to predict trait values. We improve the performance and predictive capacity thanks to the inclusion of epistatic interactions, moving beyond single-position analyses to capture the collective influence of interacting residues, and incorporation of position-specific amino acid clusters. The resulting approach remains simple, fully interpretable, and applicable with as few as one hundred known sequence-phenotype pairs. This enables both the identification of mechanistically informative positions and prediction of quantitative phenotypes for previously uncharacterized protein variants.

## Materials and Methods

### Sequence and phenotypes

Dataset compilation, sequence retrieval, alignment, and trimming procedures follow the framework described in our previous publication [25]. The thirteen studied phenotypes include the optimal growth temperature (OGT) of organisms using sequences from seven proteins: adenylate kinase (AdK) pyruvate kinase (Pk), phosphoglycerate kinase (PgK), transcription factors SPT4 and SPT5, elongation factor Tu (EF-Tu), and β-glycosidase [26]. For this last one, we also included the optimal function temperature (OFT) [26]. Additionally, we included the absorption wavelength of microbial rhodopsins [27], the maximal muscle concentrations of mammalian myoglobins [28], and percent of inhibition of HIV protease by atazanavir (ATV), saquinavir (SQV) and fosamprenavir (FPV) [29]. To compare our findings with what was already reported we conducted a literature search of relevant experimental studies for each dataset and compiled the positions reported to be relevant for phenotype modulation (Supplementary file 1).

### Identification of phenotype-modulating positions

We employed a Monte Carlo approach with Metropolis Hasting criterion to identify a subset of positions that strongly correlate to phenotypic variation. We initialized the search with a random subset of positions comprising 25% of a multiple sequence alignment (MSA). We computed the pairwise sequence Hamming distances across the selected positions (0 if the same amino acid, otherwise 1) and the absolute value of the phenotype distances. Next, we evaluated the correlation between Hamming and phenotype distances using the Spearman correlation coefficient (ρ). The subset is modified by randomly adding or removing a single position and evaluating the improvement of ρ using a Monte Carlo approach [30]. To avoid addition of poorly correlated positions we included a Lasso regularization scheme that penalizes the actual ρ (Equation (1)).

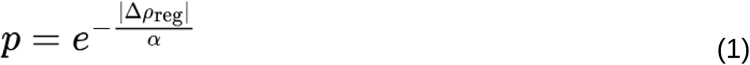

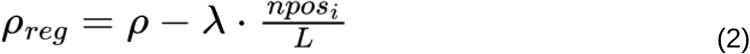

Where ρ_reg_ incorporates the L1 penalty on the subset size to prevent overfitting (Equation 2). ρ is the raw Spearman correlation obtained for the proposed subset, npos_i_ the selected positions at iteration i, and L the total number of alignment positions. The adaptive parameter α increases when no proposal was accepted for 5 consecutive iterations and reset to 1 whenever a proposal was accepted, mimicking a simulated annealing approach [31]. This protocol is applied until the subset of positions remains unchanged for 40000 iterations. The regularization parameter λ was selected using a multiobjective optimization protocol [32] to balance the raw Spearman correlation and the number of selected sequence positions (Figure S1).

### Representation of epistatic interactions in the sequences

We included epistasis by calculating the distance between two sequences using residue pairs instead of single positions. In this case, the pair contributed to the Hamming distance only when both positions differed in the compared sequences (joint-divergence rule) [33].

### Amino acid residues clustering

We implemented two algorithm variants where the distance between two sequences is calculated with a simplified sequence representation using amino acid clusters. In these cases, two amino acid residues are considered different when they belong to different clusters.The search for the optimal grouping begins at each position with 21 clusters (including gaps) and the calculation of the Spearman correlation coefficient between sequence and phenotype distances. In each subsequent step, a new grouping proposal is generated by either splitting a randomly selected group that contains multiple amino acids or merging two randomly selected groups, with equal probability. A new Spearman correlation coefficient between sequence and phenotype distances is calculated. The new grouping is then accepted or rejected using the Metropolis-Hastings criterion (Equation (3)):

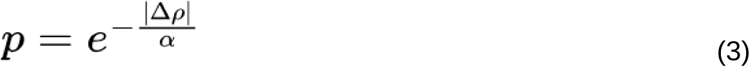

where α is an adaptive parameter that decreases every 100 iterations [31]. The optimization runs for a total of 1000 iterations per column.

Once we defined the groups, we calculated the amino acid co-occurrence Jaccard distance matrix. Briefly, taken all the thirteen cases together, we counted the times that two amino acids appear in the same group (including gaps) defining a 21 x 21 matrix:

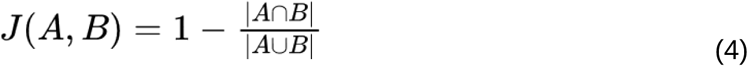

We applied Principal Component Analysis (PCA) to the amino acid co-occurrence Jaccard distance matrix to obtain a low-dimensional representation of the similarity relationships among amino acids. In this representation, amino acids with similar co-occurrence patterns across grouping schemes are positioned closer together, while dissimilar amino acids are separated along the principal components. The first principal component (PC1) captures the largest source of variation in the distance matrix, followed by the second principal component (PC2), and so on [35]. We evaluated the number of principal components to be considered using the Scree Plot [36]. We then utilized this spatial mapping of amino acids to separate them into k predefined clusters using the k-means algorithm. The optimal number of clusters (k) was determined via a Silhouette score analysis [37]. The calculations were executed by using the clustering and dimensionality reduction modules of scikit-learn library [38].

### Phenotype prediction

Phenotype prediction using the positions selected by the method was evaluated as described in our previous publication (Figure 1) [25]. Briefly, each MSA was split into training and test sets. The sequence and phenotype distances were calculated for each sequence pair in the training set, taking into account only selected positions, and a linear regression model was then fitted.

**Figure 1.**
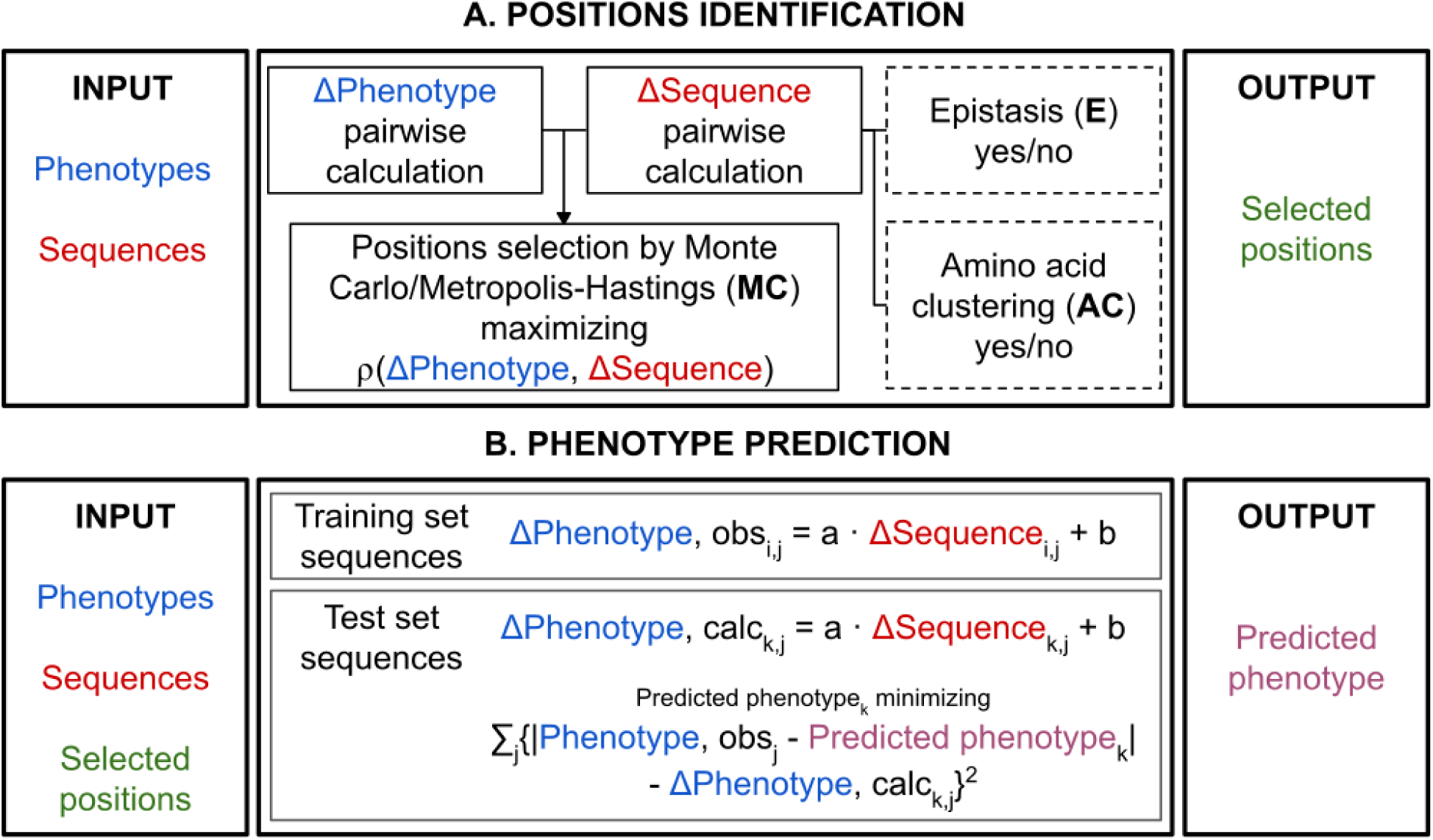
Workflow for the identification of phenotype-modulating positions and phenotype prediction. **A**. Selection of a set of positions. Based on a protein MSA where each sequence is associated with a phenotype value, the algorithm iteratively searches positions using a MC/M-H protocol for adding or removing single positions that maximize the Spearman correlation (ρ) between pairwise sequence Hamming distances and absolute phenotype distances. Position-specific amino acid clustering can be applied using an independent MC/M-H protocol that merges/splits residue groups. Second-order epistasis can also be included by evaluating distances on residue pairs. The set of single positions or position pairs with the highest ρ is reported. **B**. Phenotype prediction. The algorithm first computes a linear regression model for the correlation between pairwise sequence and phenotype distances in the training set of sequences, using the selected positions. The model is then used to predict the phenotypes for sequences from the validation set by minimizing the sum of squared differences between observed and calculated phenotype distances (see Methods). Predicted phenotypes for the validation sequences are reported.

For each test sequence, the sequence distance to all training MSA sequences was calculated. These values were used as input in the linear regression model to calculate phenotype distances from the test sequence to all test MSA sequences. The value for the phenotype of a test sequence was chosen as the one that minimizes the sum of squared differences between (1) the phenotype distances from the test sequence to all training MSA sequences calculated with the linear regression model and (2) the phenotype distance between the phenotype of a test sequence and each known phenotype from the training set. Model performance was evaluated using the R² coefficient of the correlation between the phenotypes calculated for the test sequences and their known phenotypes.

### Multi-Objective Optimization and Pareto Front Analysis

Multi-objective optimization addresses problems in which competing objectives must be balanced simultaneously, such that improving one objective may worsen another. We formulated model selection as a two-objective optimization problem with two competing goals: (i) minimizing the percentage of alignment positions selected, and (ii) maximizing performance (ρ or R^2^). The Pareto front was subsequently used to identify optimal solutions under these trade-offs. Specifically, a workflow variant A was considered to dominate a workflow variant B if (ρA >= ρB) and (A positions <= B positions), where at least one of these inequalities was strict: (ρA > ρB) or (A positions < B positions). The Pareto front was defined as the set of non-dominated variants, solutions for which no alternative variant achieves both a lower (or equal) percentage of selected positions and a higher (or equal) correlation simultaneously. Variants dominated for at least one other are excluded from the front. The Pareto front captures the optimal trade-off between model sparsity and explanatory performance [32].

“Single-point” receiver operating characteristic (ROC) analysis

The positive set (P) was defined as the set of literature-annotated residues, and the negative set (N) was defined as all positions that did not appear as positive in literature. For a given set of positions selected by our algorithm, the ones belonging to P are true positives (TP), the ones belonging to N are False Positives (FP). Similarly, the positions in P that were not selected by our algorithm are False Negatives (FN), the positions in N that were not selected are True Negatives (TN). Sensitivity and specificity are calculated using TP, FP, FN and TN values.

### Radial distribution of the selected positions

For each selected position, we compute the minimum C_α_-C_α_ distance to any functional site residue (r). Catalytic sites were retrieved from the Catalytic Site Atlas [34]. For non-enzymes, active sites or critical contact surfaces were retrieved from PDBe-KB [35]. A normalized contact distribution function (g(r)_n_) was calculated as the binned contact density for each dataset divided by the corresponding binned density of all chain contacts, normalized by the area under the curve.

## Results

### Inclusion of epistasis and amino acid clustering in the detection of phenotype-modulating positions

We build on the heuristic developed in previous work [25] by including epistasis and amino acid clustering. All variants of the algorithm take as input a multiple sequence alignment and the quantitative phenotypes associated with each sequence (Figure 1A, left). Briefly, the algorithm starts the identification of phenotype-modulating positions by calculating for each MSA position the Spearman correlation between all pairwise sequence distances and absolute phenotype distances (Figure 1A, center). The results are then used to select a set of positions that optimize the correlation between sequence and phenotype distances (Figure 1A, right). Our previous algorithm, designated here as Single Site (SS) used a greedy heuristic to select the set of positions [25]. Here, we introduce the algorithm variant Monte Carlo-Single Site (MC-SS), which evaluates combinations of positions by Monte Carlo sampling guided by a Metropolis-Hastings acceptance criterion.

We introduce the use of second-order amino acid epistasis and position-specific amino acid clustering on top of the MC-SS variant (Figure 1A, center). In the algorithm variant Monte Carlo-Epistasis (MC-E) two alignment positions are combined as a single entity and contribute to the pairwise Hamming distance only when both positions differ simultaneously between the compared sequences (see Methods). In the algorithm variants Amino Acid Clustering-Monte Carlo-Single Site (AC-MC-SS) and Amino Acid Clustering-Monte Carlo-Epistasis (AC-MC-E), amino acid residues are grouped to maximize the Spearman correlation between the Hamming and phenotype distances (see Methods). This yields a total of four new algorithm variants in addition to the published one.

The positions selected by each algorithm variant are then evaluated based on their ability to predict quantitative phenotypes in uncharacterized sequences (Figure 1B and Methods). Our original version of the algorithm was developed using 4 case studies. Here, we used 13 case studies with a wide diversity of quantitative phenotypes, including the optimal growth temperature (OGT) of four enzymes and three transcription factors, the optimal function temperature (OFT) of an enzyme, the inhibition of HIV-1 protease by three antiretroviral drugs, the skeletal muscle concentration of myoglobin across mammalian species with varying diving capacities and the retinal absorption wavelength of microbial rhodopsins (see Methods). Additionally, the sequence positions reported in the literature to be associated with each phenotype (see Methods) serve as a primary reference to evaluate the physical nature and spatial distribution of our selected positions.

### Epistasis and amino acid clustering improve the identification of phenotype modulating positions

We evaluated the global performance of five variants of the algorithm (SS, MC-SS, MC-E, AC-MC-SS and AC-MC-E) across the full set of 13 case studies. Figure 2 provides a case-by-case look for both training and prediction performance. Figure 2A compares the results of training, evaluated using the Spearman correlation coefficient (ρ) between sequence distance and phenotype distance for the set of selected positions. Substituting the greedy search of the SS algorithm (grey) for the Monte Carlo search of the MC-SS algorithm (red) improves performance in training. Introduction of epistasis in the MC-E variant (blue) and introduction of amino acid clustering in the AC-MC-SS variant (green) both improve performance relative to MC-SS. Finally, the simultaneous use of Monte Carlo search, epistasis and amino acid clustering in the AC-MC-E variant (pink) leads to the best results in training performance. Figure 2B compares the performance of the five algorithm variants in a prediction setting. The patterns are similar to those observed in Figure 2A. The simultaneous use of Monte Carlo search, epistasis and amino acid clustering in the AC-MC-E variant (pink) leads to the best overall predictions and largely outperforms the baseline SS model (grey) in most cases. We also evaluated the ability of the positions identified in the literature as important for phenotype modulation (orange). The MC-E (blue), AS-MC-SS (green) and AC-MC-E variants (pink) outperform the set of positions from the literature (orange) in most cases.

**Figure 2.**
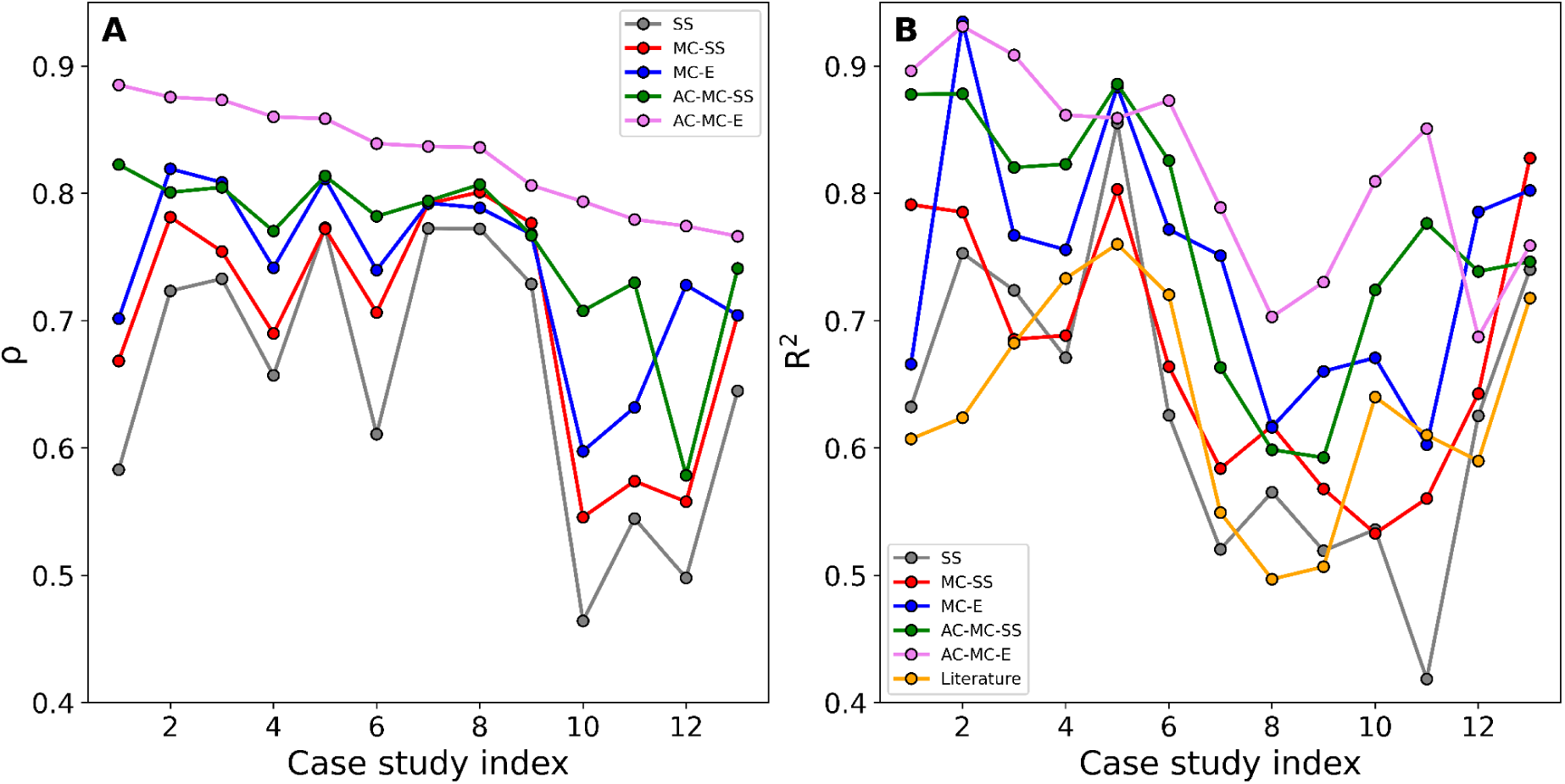
Performance of algorithm variants. **A**. Training results. Spearman correlation (ρ) between sequence distance and phenotype distance for the set of selected positions, sorted by the value for the AC-MC-E method. **B**. Prediction performance. Coefficient of determination (R^2^) for observed versus predicted phenotypes for each case study. Case studies are sorted as in panel A.

We have also compared the prediction performance of the four new algorithm variants relative to the SS variant, normalized by the maximum improvement from the SS variant. For each case study and variant we calculate R^2^_normalized_

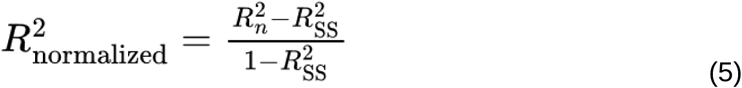

where the subindex SS refers to the SS model and the subindex n refers to variant n. The original SS variant has by definition an R^2^_normalized_ of 0. Figure 3A summarizes the averaged results for each algorithm variant as a notched box plot, where the notches show the 95% confidence interval of the median. Incorporation of the Monte Carlo search in the MC-SS algorithm (red) yields a statistically significant improvement (p-value 0.05) in prediction performance. The MC-SS variant is further improved to similar degrees by incorporation of epistasis in the MC-E variant (blue) or incorporation of amino acid clustering in the AC-MC-SS variant (green). The AC-MC-E variant (pink) using all of Monte Carlo search, epistasis and amino acid clustering presents the best prediction performance. The prediction performance of the positions identified in the literature as important for phenotype modulation is much lower and comparable to that of the original SS algorithm (orange).

**Figure 3.**
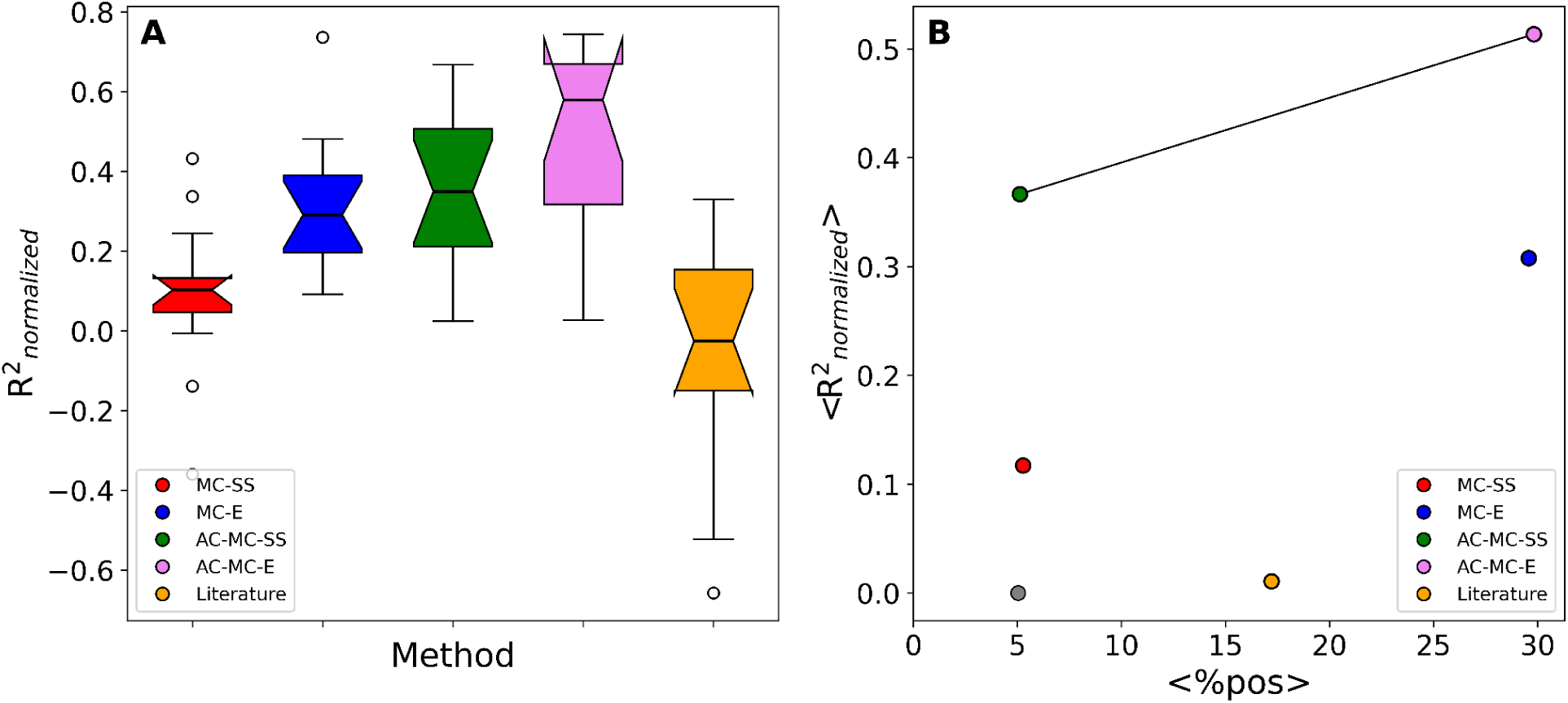
Predictive performance of algorithm variants relative to the SS variant. **A**. Box-plot of the distribution of the fraction of improvement in R^2^ relative to the SS variant across case studies. Notches show the 95% confidence interval of the median. **B**. Multi-objective optimization analysis. Percentage of alignment positions selected by the algorithm against the average fraction of R^2^ improvement. Solutions dominated in at least one objective are excluded from the Pareto front, which is highlighted by a black line.

We also evaluated the relationship between the improvement in the prediction performance and the fraction of alignment positions selected by the algorithm. An ideal algorithm would have both a high prediction performance and a low percentage of selected positions. The Pareto front in Figure 3B considers both aspects of this multi-objective optimization problem. In this framework, we consider that an algorithm variant solution A outperforms B if A is no worse than B in both objectives, and better than B in at least one objective (see Methods). Predictions using the positions identified in the literature as important for phenotype modulation are dominated by one or more algorithm variants. On the other hand, algorithm variants AC-MC-SS and AC-MC-E are both top-performing solutions to the problem and constitute the Pareto front. Consequently, we focus the following sections on the analysis of these two variants.

### Amino acid clustering reveals a role of hydrophobicity in phenotype modulation

Results in the previous section show that amino acid clustering improves the prediction of phenotype modulating positions. The clusters defined by the algorithm are those that optimize the sequence-phenotype relationship at each MSA position of the 13 study cases. We have investigated the average properties of these clusters to better understand the basis for the improvement in the prediction performance. We first calculated amino acid co-occurrence Jaccard distance matrices for the AC-MC-SS and AC-MC-E algorithm variants (see Methods). Next, we calculated the principal components (PCs) of each co-occurrence matrix (see Methods). We found that the combination of the first and second components captured more than 40% of the total variance for both AC-MC-SS and AC-MC-E, with additional components capturing minor fractions of the variance. The results are shown in Figure 4. The first principal component shows a strong correlation with amino acid hydrophobicity (R2 0.61 and 0.62 for AC-MC-SS and AC-MC-E variants, using the normalized hydrophobicity scales reported in Cid et al.[36] Figure S9). This is in agreement with hydrophobicity being a major contributor to amino acid substitution evolution in proteins [37,38]. Conversely, we were not able to find a statistically significant correlation between the second principal component over 566 amino acid properties [39], suggesting that this component is a complex mixture of multiple contributions.

**Figure 4.**
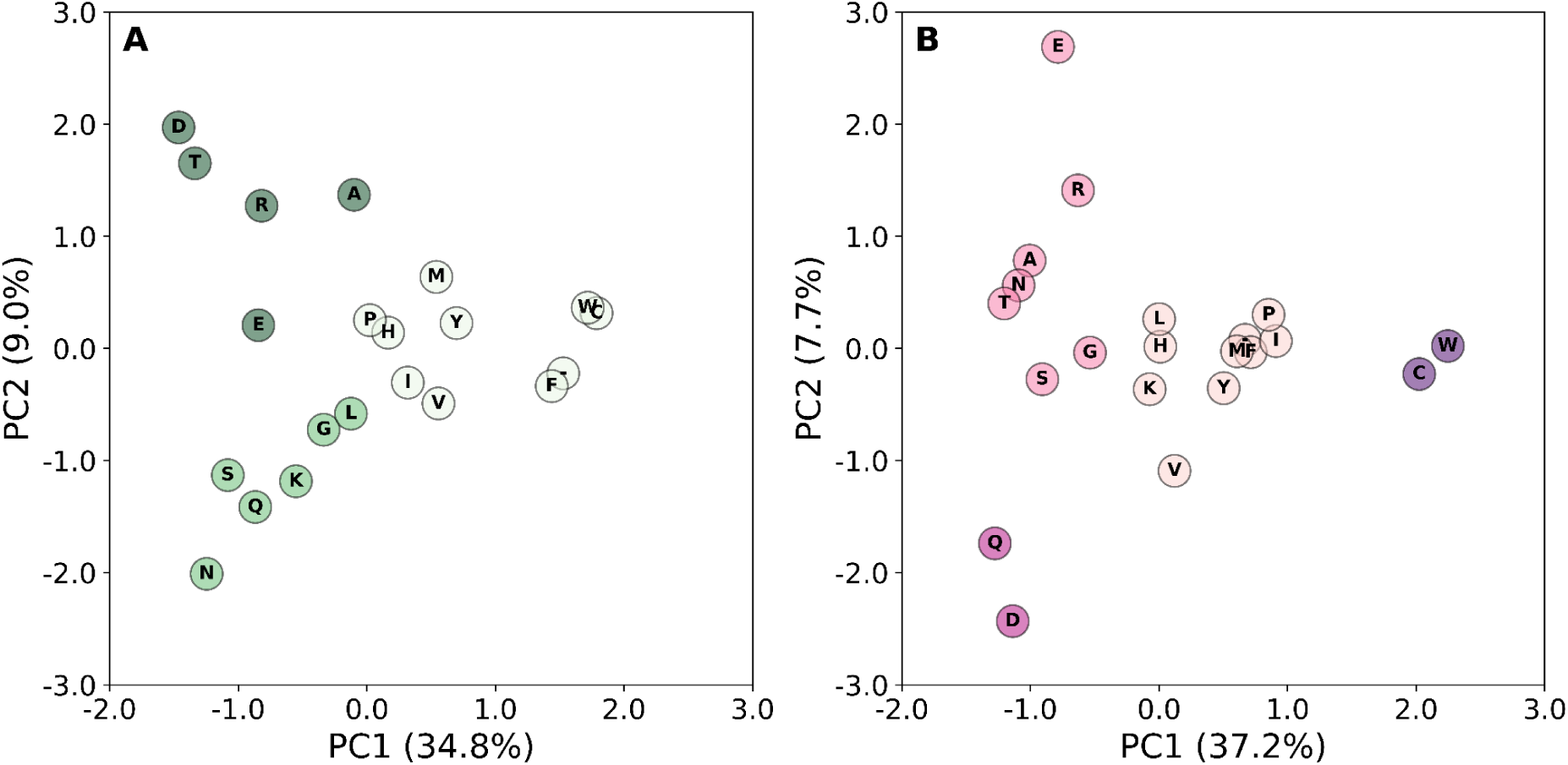
Averaged structure of the amino acid clustering performed by our algorithm. Two-dimensional projection of the first two principal components calculated from the Jaccard distance matrix of amino acid co-occurrence in all MSA positions of the 13 study cases. Axis labels indicate the percentage of total variance explained by the first (PC1) and second (PC2) principal components, respectively. Clusters were partitioned using the K-means algorithm, where the optimal number of clusters (k) was determined by maximizing the average silhouette score across a range of values. Each color represents a cluster. **A**. AC-MC-SS. **B**. AC-MC-E.

The vast majority of variability is effectively captured by 3 and 4 groups for Figure 4A and 4B, respectively, suggesting a small effective size for the amino acid alphabet capturing the modulation of quantitative phenotypes. On the other hand, we could not find a complete rationale for the composition of each cluster. Although it seems quite clear that hydrophobicity guides clusterization, this property does not fully rationalize our results. In all, the averaged amino acid clustering performed by the AC-MC-SS and AC-MC-E algorithm variants presents both intuitive and more intricate combinations of physicochemical features.

### Single-site and epistasis analyses identify phenotype modulating positions at different distances from the protein functional sites

Our Pareto front analysis selects two top-performing algorithm variants. Variant AC-MC-SS yields a good prediction performance using an average of 5% of positions in the alignment. Variant AC-MC-E yields a better prediction performance but uses an average of 30% of alignment positions. Visual inspection of the selected positions in the 13 study cases revealed a pattern. Figure 5 shows a representative example, the percent Inhibition of HIV-1 protease by FPV. The AC-MC-SS algorithm variant is able to predict inhibition (R2 = 0.60) using only 10 sequence positions, most of which are close to the inhibitor binding site. In contrast with this, the AC-MC-E algorithm variant predicts inhibition (R2 = 0.70) using 40 sequence positions. These sequence positions are broadly distributed throughout the protein scaffold.

**Figure 5.**
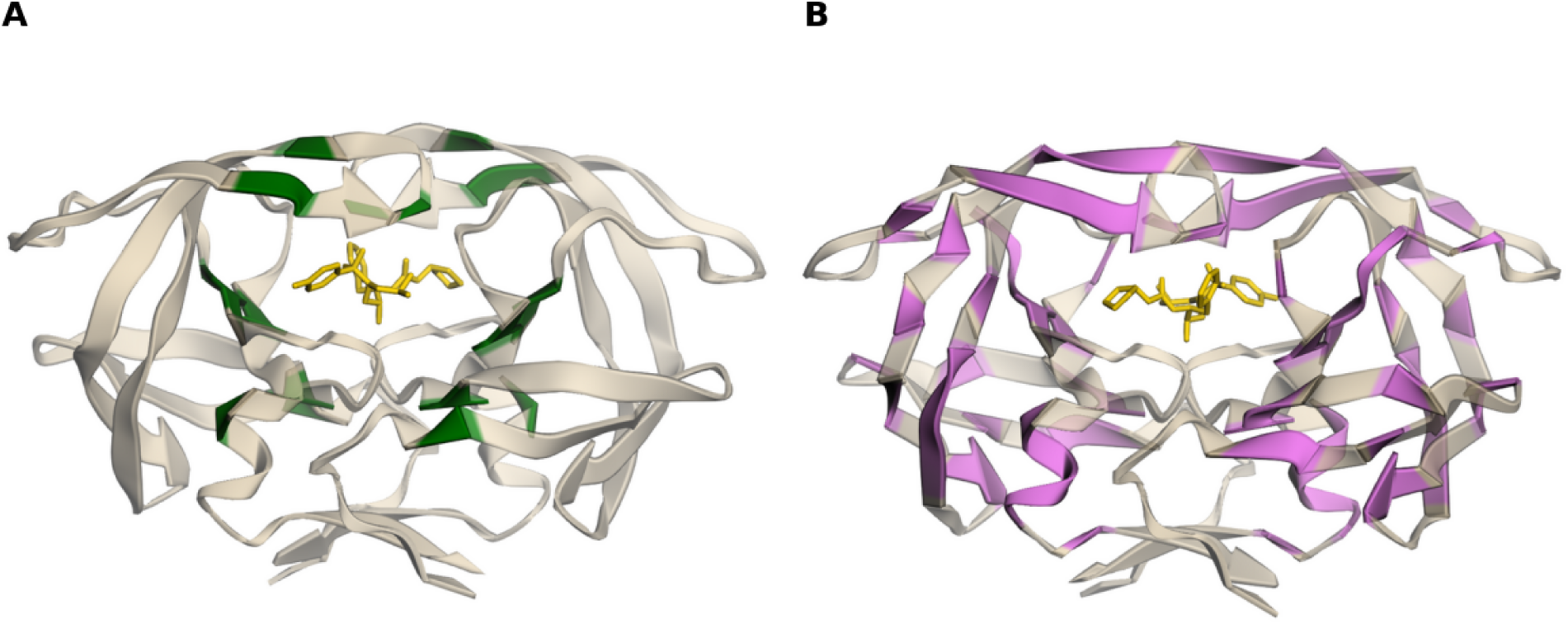
Percent Inhibition of HIV-1 protease by FPV. **A**. Distribution of positions selected by the AC-MC-SS algorithm variant in the HIV-1 protease (PDB:3ekv). The active metabolite of FPV is shown in yellow. **B**. Distribution of positions selected by the AC-MC-E algorithm variant in the HIV-1 protease (PDB:3ekv). The active metabolite of FPV is shown in yellow.

We evaluated the spatial distribution of the positions selected by the AC-MC-SS and AC-MC-E algorithm variants by calculating the distance relative to catalytic sites for enzymes and active sites or critical contact surfaces for non-enzymes, collectively called “functional sites” (g(r)n). As shown in Figure 6, the distance distribution for the AC-MC-SS method shows a peak at about 8 Å (continuous green line). In contrast, the AC-MC-E algorithm variant displays a peak of lower height close to 8 Å and a wider distance distribution (continuous pink line). This closely resembles the random control distributions generated by selecting the same number of positions as the AC-MC-SS and AC-MC-E algorithm variants (dashed green and pink lines, respectively). These control distributions show a poor prediction performance (Figure S5). This result suggests that modulation of quantitative phenotypes involves positions proximal and distal from the protein functional sites in our 13 study cases. Phenotype modulation by proximal sites can be captured by studying single sites, while phenotype modulation by distal sites involves epistasis and can be captured by considering pairs of residues.

**Figure 6.**
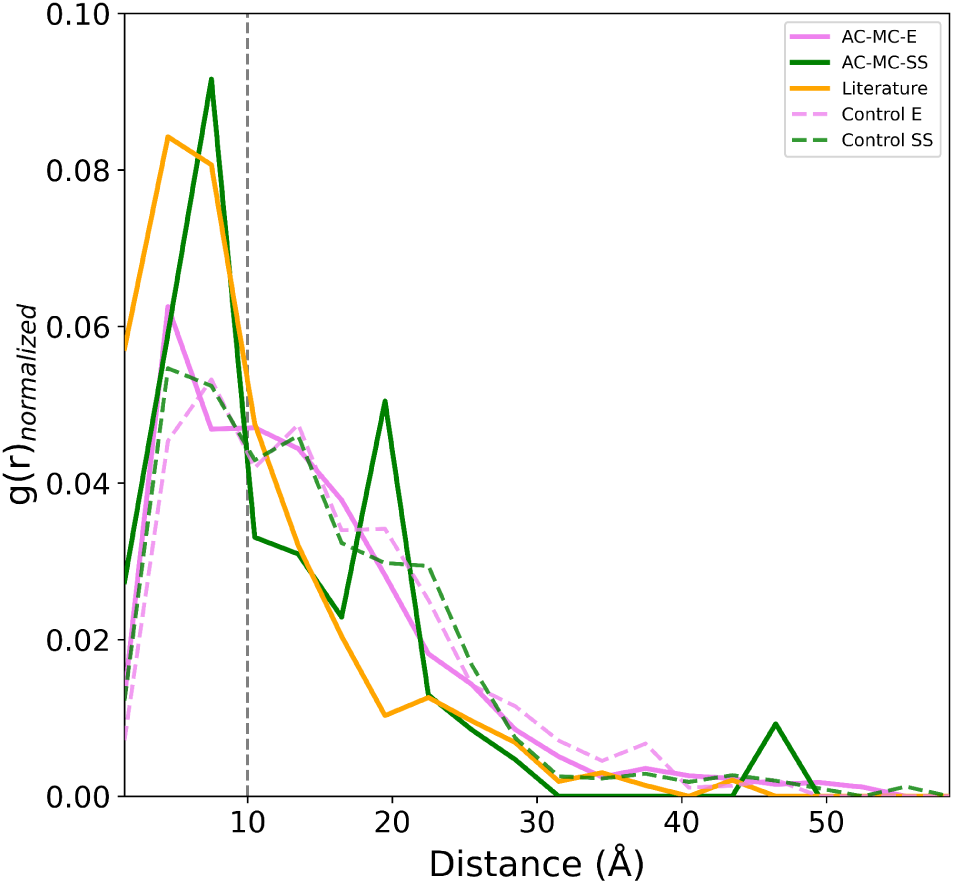
Distance distribution of positions selected by the AC-MC-SS and AC-MC-E algorithm variants, relative to the protein functional sites. Solid pink and green curves represent <g(r)n> calculations for AC-MC-E and AC-MC-SS selected positions respectively. Controls were generated by using an equivalent number of randomly selected positions or position pairs (dashed pink and green lines). The solid orange curve represents literature selected positions.

### Comparison to phenotype-modulating positions identified in the literature

The AC-MC-SS and AC-MC-E algorithm variants. outperform the set of positions from the literature (orange) in the prediction of quantitative phenotypes (Figures 2 and 3). This somewhat unexpected result prompted us to study the overlap between the positions reported in the literature and those selected by the AC-MC-SS and AC-MC-E algorithm variants. The Jaccard index for the AC-MC-SS and AC-MC-E algorithm variants has median values of 0.05 and 0.12, respectively (Figure 7A).

**Figure 7.**
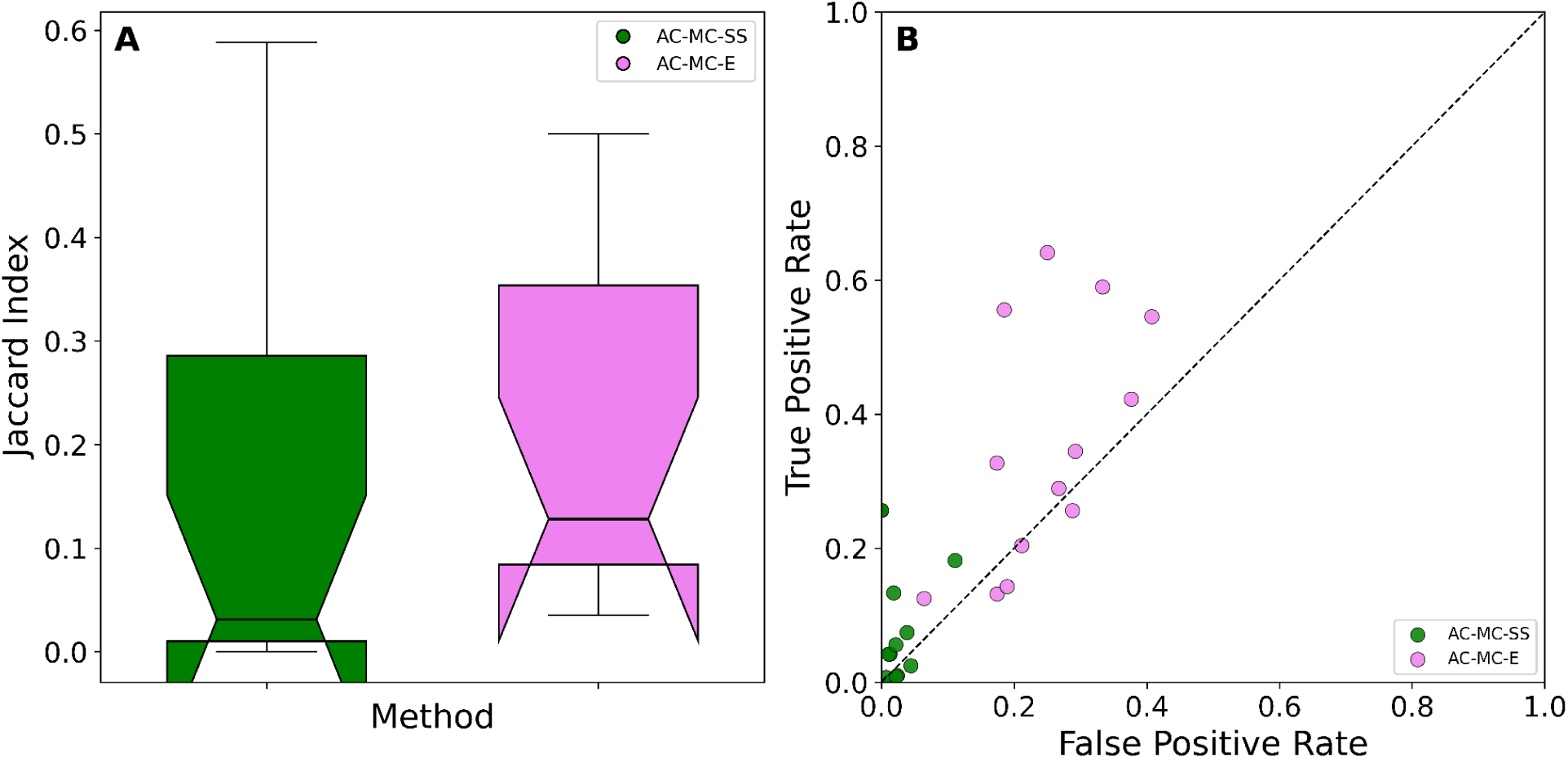
Comparison to phenotype-modulating positions identified in the literature. **A**. Box-plot of the distribution of the Jaccard index for the overlap between positions identified in the literature and positions selected by the AC-MC-SS and AC-MC-E algorithm variants. Notches show the 95% confidence interval of the median. **B**. Performance of the AC-MC-SS and AC-MC-E algorithm variants as categorical predictors of the positions identified in the literature. Each point represents a single case study. Green circles: AC-MC-SS algorithm variant. Pink circles: AC-MC-E algorithm variant. The dashed line represents a random predictor (TPR = FPR).

We have also evaluated the performance of our algorithm variants as categorical predictors of the positions reported in the literature. Since our algorithm variants do not have a variable threshold that can be used to generate a receiver-operating characteristic (ROC) curve, the results are single-points in the ROC space (Figure 7B). For each case study and algorithm variant, we calculated the sensitivity (True positive rate, TPR) and specificity (False positive rate, FPR) of the selected positions. Most of the case studies lie near the diagonal line expected for a random prediction, where TPR = FPR. In short, our algorithm is a poor predictor of annotated literature positions. A notable exception are the three HIV-1 protease study cases, where resistance-determining positions have been exhaustively classified in the literature and the distance to the diagonal is higher (HIV-FPV: TRP=0.64, FRP=0.25; HIV-SQV: TRP=0.56, FRP=0.19; HIV-ATV: TRP=0.59, FRP=0.33). This, together with the low degree of overlap shown in Figure 7A, suggests that identification of phenotype-modulating positions in the literature is far from complete and that our algorithm can contribute to this task.

## Discussion

To contextualize the performance of the proposed strategy, we benchmarked it against phylogenetic imputation of continuous traits under a Brownian motion model [40] and an ESM-2 embedding-based predictor using ridge regression [21] across our thirteen datasets, which span distinct molecular and evolutionary regimes. While these methodologies represent state-of-the-art algorithms for decoding functional protein characteristics from sequence information, phylogenetic imputation does this by taking into account phylogenetic history, while the ESM-2 embeddings consider contextual sequence information without explicitly considering phylogenetic information. The comparative performance can be seen in **Figure 8A**. In the eight datasets where the phenotype is a temperature (ADK, PK, PGK, SPT4, SPT5, EFTU, GH1-OGT and GH1-OFT), the variants incorporating amino acid alphabet reduction and Monte Carlo search achieved the highest R^2^ in seven cases. This overall pattern suggests that the proposed strategy effectively exploits comparative information in ortholog alignments to predict phenotypes that are temperatures. The contrast between the two GH1 datasets is particularly informative: all methods performed better when predicting the organism’s optimal growth temperature (OGT) than the enzyme’s optimal functional temperature (OFT), despite both tasks relying on the same alignment. The marked reduction in ESM-2 performance for OFT (R^2^=0.24), alongside the strong performance of AC-MC-E (R^2^=0.81), illustrates that sequence embeddings do not necessarily capture the determinants of a specific molecular phenotype as effectively as approaches that exploit variation across homologous sequences. In the three HIV-1 protease resistance datasets, AC-MC-E achieved the highest R^2^ in every case, consistent with the relevance of epistatic interactions in resistance evolution [41,42] and the limitations of phylogenetic imputation for traits subject to convergent selection. Conversely, ESM-2 performed best for the maximal absorption wavelength of bacteriorhodopsins with R^2^=0.87, while AC-MC-E achieved a lower but also comparable performance of R^2^=0.76. This result is consistent with the possibility that spectral tuning, which depends strongly on a limited number of residues surrounding the retinal chromophore, is better captured by representations that encode local sequence context than by approaches relying on comparative patterns across homologs. For muscular myoglobin concentration in diving mammals (Mb), phylogenetic imputation achieved the highest R^2^ followed by AC-MC-SS. The relatively high sequence similarity within this group may support more reliable phylogenetic reconstruction and make the detection of epistasis by the AC-MC-E algorithm variant more difficult.

**Figure 8.**
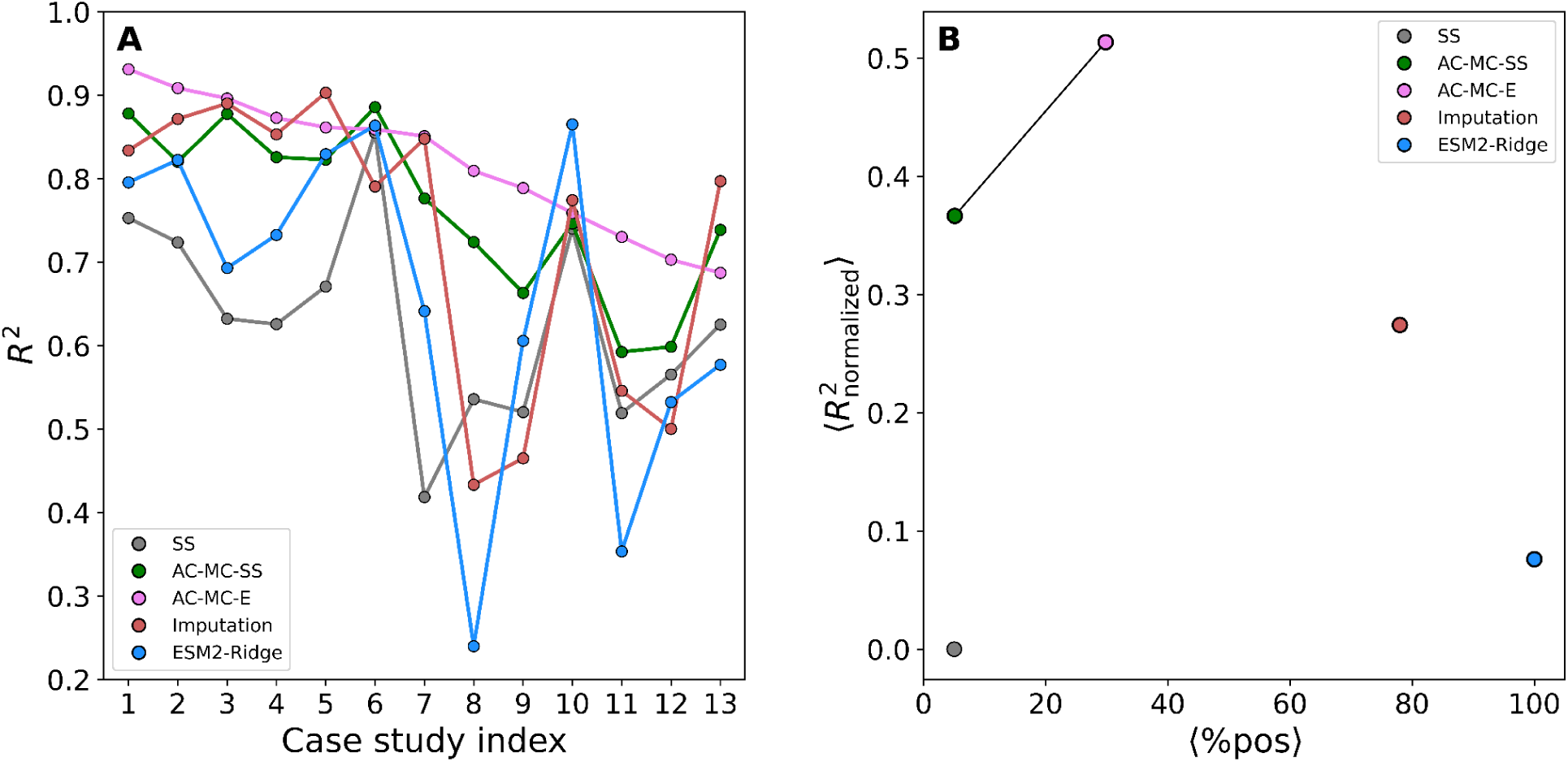
Comparative algorithm variants performance. **A**. Prediction performance. Coefficient of determination (R^2^) for observed versus predicted phenotypes for each case study. Case studies are sorted by the values of the AC-MC-E variant. **B**. Multi-objective optimization analysis. Percentage of alignment positions selected by the algorithms against the average fraction of R^2^ improvement. Solutions dominated in at least one objective are excluded from the Pareto front, which is highlighted by a black line.

As it is seen in **Figure 8A**, the proposed strategy, particularly AC-MC-E, displayed a comparatively uniform performance profile: it ranked among the top three methods in all thirteen datasets and achieved the highest R^2^ in nine. Neither AC-MC-E nor AC-MC-SS ranked last in any dataset, and the lowest R^2^ obtained by AC-MC-E (0.687 for Mb) exceeded the corresponding minima of the other four methods. Furthermore, AC-MC-E achieves a higher predictive performance than AC-MC-SS in 11 of the 13 cases, highlighting the relevance of including epistatic interactions in the model. Interestingly, **Figure 8B** also shows that both phylogenetics and machine learning methods are far from the Pareto front, both outperformed by AC-MC-SS and AC-MC-E, which achieved a better predictive performance while using significantly lower percentages of alignment positions. These results suggest that combining amino acid alphabet reduction with Monte Carlo search provides a useful balance between predictive performance and robustness across heterogeneous sequence–phenotype relationships. The additional advantage of the epistatic layer in the HIV-1 resistance datasets further suggests that the framework can accommodate phenotypes for which the effects of individual substitutions depend on their sequence context. Within this scope, the findings support the value of exploiting homologous sequence variation and, where appropriate, epistatic structure to predict quantitative phenotypes, while highlighting that the most suitable approach depends on the biological and evolutionary determinants of the property under study.

## Acknowledgments

We thank Dr. Diego U. Ferreiro (Universidad de Buenos Aires) and Ezequiel A. Galpern (Centre for Genomic Regulation) for carefully reading and discussing this manuscript. Finally, we manifest our deep concerns about the sudden discontinuation of scientific funding which impedes the development of new projects and terminates the current ones.

In loving memory of Miguel A. Roman.

## Abbreviations

E: Epistatic
SS: Single site
MC: Monte Carlo
M-H: Metropolis-Hastings
AC: Amino acid clustering
OGT: Optimal Growth Temperature
OFT: Optimal Function Temperature
MSA: Multiple Sequence Alignment
Adk: Adenylate kinase
Pk: Pyruvate kinase
Pgk: Phosphoglycerate kinase
SPT4: Transcription factor SPT4
SPT5: Transcription factor SPT5
Ef-Tu: Elongation factor Tu
IP: Inhibitory potency
BR: microbial rhodopsin
MB: Myoglobin
HIV-PR: HIV-1 protease
SQV: saquinavir
FQV: fosamprenavir
ATV: atazanavir
GH1: *β*-glycosilase hydrolase 1
AUC: area under the curve
ROC: Receiver Operating Characteristic
TPR: True Positive Rate
FPR: False Positive Rate.

## References

[1] M. Baek, F. DiMaio, I. Anishchenko, J. Dauparas, S. Ovchinnikov, G.R. Lee, J. Wang, Q. Cong, L.N. Kinch, R.D. Schaeffer, C. Millán, H. Park, C. Adams, C.R. Glassman, A. DeGiovanni, J.H. Pereira, A.V. Rodrigues, A.A. van Dijk, A.C. Ebrecht, D.J. Opperman, T. Sagmeister, C. Buhlheller, T. Pavkov-Keller, M.K. Rathinaswamy, U. Dalwadi, C.K. Yip, J.E. Burke, K.C. Garcia, N.V. Grishin, P.D. Adams, R.J. Read, D. Baker, Accurate prediction of protein structures and interactions using a three-track neural network, Science 373 (2021) 871–876.

[2] J. Jumper, R. Evans, A. Pritzel, T. Green, M. Figurnov, O. Ronneberger, K. Tunyasuvunakool, R. Bates, A. Žídek, A. Potapenko, A. Bridgland, C. Meyer, S.A.A. Kohl, A.J. Ballard, A. Cowie, B. Romera-Paredes, S. Nikolov, R. Jain, J. Adler, T. Back, S. Petersen, D. Reiman, E. Clancy, M. Zielinski, M. Steinegger, M. Pacholska, T. Berghammer, S. Bodenstein, D. Silver, O. Vinyals, A.W. Senior, K. Kavukcuoglu, P. Kohli, D. Hassabis, Highly accurate protein structure prediction with AlphaFold, Nature 596 (2021) 583–589.

[3] C.L. Araya, D.M. Fowler, Deep mutational scanning: assessing protein function on a massive scale, Trends Biotechnol. 29 (2011) 435–442.

[4] D.M. Fowler, S. Fields, Deep mutational scanning: a new style of protein science, Nat. Methods 11 (2014) 801–807.

[5] J. Pei, N.V. Grishin, AL2CO: calculation of positional conservation in a protein sequence alignment, Bioinformatics 17 (2001) 700–712.

[6] E. Teppa, A.D. Wilkins, M. Nielsen, C.M. Buslje, Disentangling evolutionary signals: conservation, specificity determining positions and coevolution. Implication for catalytic residue prediction, BMC Bioinformatics 13 (2012) 235.

[7] J.R. Manning, E.R. Jefferson, G.J. Barton, The contrasting properties of conservation and correlated phylogeny in protein functional residue prediction, BMC Bioinformatics 9 (2008) 51.

[8] R.C. Lua, S.J. Wilson, D.M. Konecki, A.D. Wilkins, E. Venner, D.H. Morgan, O. Lichtarge, UET: a database of evolutionarily-predicted functional determinants of protein sequences that cluster as functional sites in protein structures, Nucleic Acids Res. 44 (2016) D308–12.

[9] O. Lichtarge, H.R. Bourne, F.E. Cohen, An evolutionary trace method defines binding surfaces common to protein families, J. Mol. Biol. 257 (1996) 342–358.

[10] T. Teşileanu, L.J. Colwell, S. Leibler, Protein sectors: statistical coupling analysis versus conservation, PLoS Comput. Biol. 11 (2015) e1004091.

[11] O. Rivoire, K.A. Reynolds, R. Ranganathan, Evolution-based functional decomposition of proteins, PLoS Comput. Biol. 12 (2016) e1004817.

[12] C.K. Cornwallis, A.S. Griffin, A guided tour of phylogenetic comparative methods for studying trait evolution, Annu. Rev. Ecol. Evol. Syst. 55 (2024) 181–204.

[13] A.E. Dewar, L.J. Belcher, S.A. West, A phylogenetic approach to comparative genomics, Nat. Rev. Genet. 26 (2025) 395–405.

[14] J. Bruggeman, J. Heringa, B.W. Brandt, PhyloPars: estimation of missing parameter values using phylogeny, Nucleic Acids Res 37 (2009) W179–84.

[15] F. Li, L. Yuan, H. Lu, G. Li, Y. Chen, M.K.M. Engqvist, E.J. Kerkhoven, J. Nielsen, Deep learning-based kcat prediction enables improved enzyme-constrained model reconstruction, Nat. Catal. 5 (2022) 662–672.

[16] J. Wang, Z. Yang, C. Chen, G. Yao, X. Wan, S. Bao, J. Ding, L. Wang, H. Jiang, MPEK: a multitask deep learning framework based on pretrained language models for enzymatic reaction kinetic parameters prediction, Brief. Bioinform. 25 (2024) bbae387.

[17] V.S. Boorla, C.D. Maranas, CatPred: a comprehensive framework for deep learning in vitro enzyme kinetic parameters, Nat. Commun. 16 (2025) 2072.

[18] S.A. Frazer, M. Baghbanzadeh, A. Rahnavard, K.A. Crandall, T.H. Oakley, Discovering genotype-phenotype relationships with machine learning and the Visual Physiology Opsin Database (VPOD), Gigascience 13 (2024) giae073.

[19] M. Karasuyama, K. Inoue, R. Nakamura, H. Kandori, I. Takeuchi, Understanding colour tuning rules and predicting absorption wavelengths of microbial rhodopsins by data-driven machine-learning approach, Sci. Rep. 8 (2018) 15580.

[20] H. Liu, G. Zhu, L. Chen, H. Ye, Y. Zhang, G. Han, Machine learning prediction of bacterial optimal growth temperature from protein domain signatures reveals thermoadaptation mechanisms, BMC Genomics 26 (2025) 957.

[21] E. Komp, K.E. Johansson, N.P. Gauthier, J.E. Gado, K. Lindorff-Larsen, G.T. Beckham, Accessible, uniform protein property prediction with a scikit-learn based toolset AIDE, Bioinformatics 41 (2025). 10.1093/bioinformatics/btaf544.

[22] L.E. Jessen, I. Hoof, O. Lund, M. Nielsen, SigniSite: Identification of residue-level genotype-phenotype correlations in protein multiple sequence alignments, Nucleic Acids Res. 41 (2013) W286–91.

[23] O.V. Kalinina, A.A. Mironov, M.S. Gelfand, A.B. Rakhmaninova, Automated selection of positions determining functional specificity of proteins by comparative analysis of orthologous groups in protein families, Protein Sci. 13 (2004) 443–456.

[24] F. Pazos, A. Rausell, A. Valencia, Phylogeny-independent detection of functional residues, Bioinformatics 22 (2006) 1440–1448.

[25] A.S. Hernández Berthet, A.A. Aptekmann, J. Tejero, I.E. Sánchez, M.E. Noguera, E.A. Roman, Associating protein sequence positions with the modulation of quantitative phenotypes, Arch. Biochem. Biophys. 755 (2024) 109979.

[26] G. Gamiz-Arco, L.I. Gutierrez-Rus, V.A. Risso, B. Ibarra-Molero, Y. Hoshino, D. Petrović, J. Justicia, J.M. Cuerva, A. Romero-Rivera, B. Seelig, J.A. Gavira, S.C.L. Kamerlin, E.A. Gaucher, J.M. Sanchez-Ruiz, Heme-binding enables allosteric modulation in an ancient TIM-barrel glycosidase, Nature Communications 12 (2021) 380.

[27] M. Karasuyama, K. Inoue, R. Nakamura, H. Kandori, I. Takeuchi, Understanding Colour Tuning Rules and Predicting Absorption Wavelengths of Microbial Rhodopsins by Data-Driven Machine-Learning Approach, Scientific Reports 8 (2018) 15580.

[28] S. Mirceta, A.V. Signore, J.M. Burns, A.R. Cossins, K.L. Campbell, M. Berenbrink, Evolution of Mammalian Diving Capacity Traced by Myoglobin Net Surface Charge, Science (2013). 10.1126/science.1234192.

[29] S.-Y. Rhee, M.J. Gonzales, R. Kantor, B.J. Betts, J. Ravela, R.W. Shafer, Human immunodeficiency virus reverse transcriptase and protease sequence database, Nucleic Acids Res 31 (2003) 298–303.

[30] X.S. Yang, 2 - Mathematical foundations, in: Introduction to Algorithms for Data Mining and Machine Learning, 2019: pp. 19–43.

[31] A. Khachaturyan, S. Semenovsovskaya, B. Vainshtein, The thermodynamic approach to the structure analysis of crystals, Acta Crystallogr. A 37 (1981) 742–754.

[32] A. Chinchuluun, P.M. Pardalos, A survey of recent developments in multiobjective optimization, Annals of Operations Research 154 (2007) 29–50.

[33] A. Biswas, A. Haldane, R.M. Levy, Limits to detecting epistasis in the fitness landscape of HIV, PLoS One 17 (2022) e0262314.

[34] A.J.M. Ribeiro, G.L. Holliday, N. Furnham, J.D. Tyzack, K. Ferris, J.M. Thornton, Mechanism and Catalytic Site Atlas (M-CSA): a database of enzyme reaction mechanisms and active sites, Nucleic Acids Res 46 (2018) D618–D623.

[35] PDBe-KB consortium, PDBe-KB: a community-driven resource for structural and functional annotations, Nucleic Acids Res 48 (2020) D344–D353.

[36] H. Cid, M. Bunster, M. Canales, F. Gazitúa, Hydrophobicity and structural classes in proteins, Protein Eng 5 (1992) 373–375.

[37] M.J. Jiménez-Santos, M. Arenas, U. Bastolla, Influence of mutation bias and hydrophobicity on the substitution rates and sequence entropies of protein evolution, PeerJ 6 (2018) e5549.

[38] A.R. Kinjo, K. Nishikawa, Eigenvalue analysis of amino acid substitution matrices reveals a sharp transition of the mode of sequence conservation in proteins, Bioinformatics 20 (2004) 2504–2508.

[39] S. Kawashima, P. Pokarowski, M. Pokarowska, A. Kolinski, T. Katayama, M. Kanehisa, AAindex: amino acid index database, progress report 2008, Nucleic Acids Res 36 (2008) D202–5.

[40] S. Guindon, J.-F. Dufayard, V. Lefort, M. Anisimova, W. Hordijk, O. Gascuel, New algorithms and methods to estimate maximum-likelihood phylogenies: assessing the performance of PhyML 3.0, Syst Biol 59 (2010) 307–321.

[41] A. Biswas, A. Haldane, E. Arnold, R.M. Levy, Epistasis and entrenchment of drug resistance in HIV-1 subtype B, Elife 8 (2019). 10.7554/eLife.50524.

[42] T.-H. Zhang, L. Dai, J.P. Barton, Y. Du, Y. Tan, W. Pang, A.K. Chakraborty, J.O. Lloyd-Smith, R. Sun, Predominance of positive epistasis among drug resistance-associated mutations in HIV-1 protease, PLoS Genet 16 (2020) e1009009.

[43] J. Castresana, Selection of conserved blocks from multiple alignments for their use in phylogenetic analysis, Mol. Biol. Evol. 17 (2000) 540–552.

[44] S. Kalyaanamoorthy, B.Q. Minh, T.K.F. Wong, A. von Haeseler, L.S. Jermiin, ModelFinder: fast model selection for accurate phylogenetic estimates, Nat. Methods 14 (2017) 587–589.

[45] B.Q. Minh, H.A. Schmidt, O. Chernomor, D. Schrempf, M.D. Woodhams, A. von Haeseler, R. Lanfear, IQ-TREE 2: New models and efficient methods for phylogenetic inference in the genomic era, Mol. Biol. Evol. 37 (2020) 1530–1534.

[46] S. Guindon, J.-F. Dufayard, V. Lefort, M. Anisimova, W. Hordijk, O. Gascuel, New algorithms and methods to estimate maximum-likelihood phylogenies: assessing the performance of PhyML 3.0, Syst. Biol. 59 (2010) 307–321.

[47] D.C. Nickle, L. Heath, M.A. Jensen, P.B. Gilbert, J.I. Mullins, S.L. Kosakovsky Pond, HIV-specific probabilistic models of protein evolution, PLoS One 2 (2007) e503.

[48] L.J. Revell, phytools 2.0: an updated R ecosystem for phylogenetic comparative methods (and other things), PeerJ 12 (2024) e16505.

[49] Z. Lin, H. Akin, R. Rao, B. Hie, Z. Zhu, W. Lu, N. Smetanin, R. Verkuil, O. Kabeli, Y. Shmueli, A. Dos Santos Costa, M. Fazel-Zarandi, T. Sercu, S. Candido, A. Rives, Evolutionary-scale prediction of atomic-level protein structure with a language model, Science 379 (2023) 1123–1130.

[50] E. Komp, K.E. Johansson, N.P. Gauthier, J.E. Gado, K. Lindorff-Larsen, G.T. Beckham, Accessible, uniform protein property prediction with a scikit-learn based toolset AIDE, Bioinformatics 41 (2025) btaf544.

