## Supplementary Information for "Amino acid clustering and epistasis guide sequence-based prediction of phenotype modulation"

### Predictive performance comparison to other existing methods

#### *Phylogenetic imputation*

For each of the thirteen studied cases, predictive performance was assessed by stratified hold-out validation: 25 sequences per case were assigned to a test set (with exception of 13 and 17 for myoglobin and GH1, respectively, which contains fewer sequences) and the remaining sequences were used for training. Alignments were trimmed with Gblocks [43] in protein mode, allowing gap positions present in a minority of sequences (-t=p -b5=h). The best-fitting amino acid substitution model was selected with ModelFinder [44] in IQ-TREE 2 [45] according to the Bayesian information criterion. Maximum-likelihood trees were *de novo* inferred with phyML 3 [46] under the selected model, with the proportion of invariable sites and the gamma shape parameter (four rate categories) estimated from the data, and midpoint-rooted. For the HIV case, the tree was inferred directly in IQ-TREE because the selected HIV model [47] is not available in phyML. Trees were inferred from all sequences, including test sequences; only the trait values of test sequences were masked, mirroring the intended use case in which the query sequence is known but its phenotype is not. Missing trait values of test-set tips were imputed with the phylo.impute function of phytools 2 [48], which fits a Brownian-motion model of trait evolution by maximum likelihood and returns maximum-likelihood estimates of missing tip values conditional on the tree and the observed data. Predictive performance on the test sets was quantified with standard accuracy and error metrics computed in R.

### Machine learning predictions using ESM-2 embeddings

We implemented a predictor based on representations from the ESM-2 protein language model [49] using the AIDE package [50]. Each sequence was represented by the mean of its per-residue embeddings (mean pooling) from the esm2_t6_8M_UR50D checkpoint (a similar performance was observed for the esm2_t33_650M_UR50D version). A ridge regression using the python package scikit-learn was fitted on these vectors, with prior standardization of the features. For the antiviral resistance cases, the response variable was $\log_{10}$-transformed and predictions were returned to the original scale ($10^x$, truncated at zero). For all other properties, the original measurement scale was used. Evaluation was performed on exactly the same holdout set described for phylogenetic imputation. Performance was quantified with the coefficient of determination ($R^2 = 1 - SS\_res/SS\_tot$).

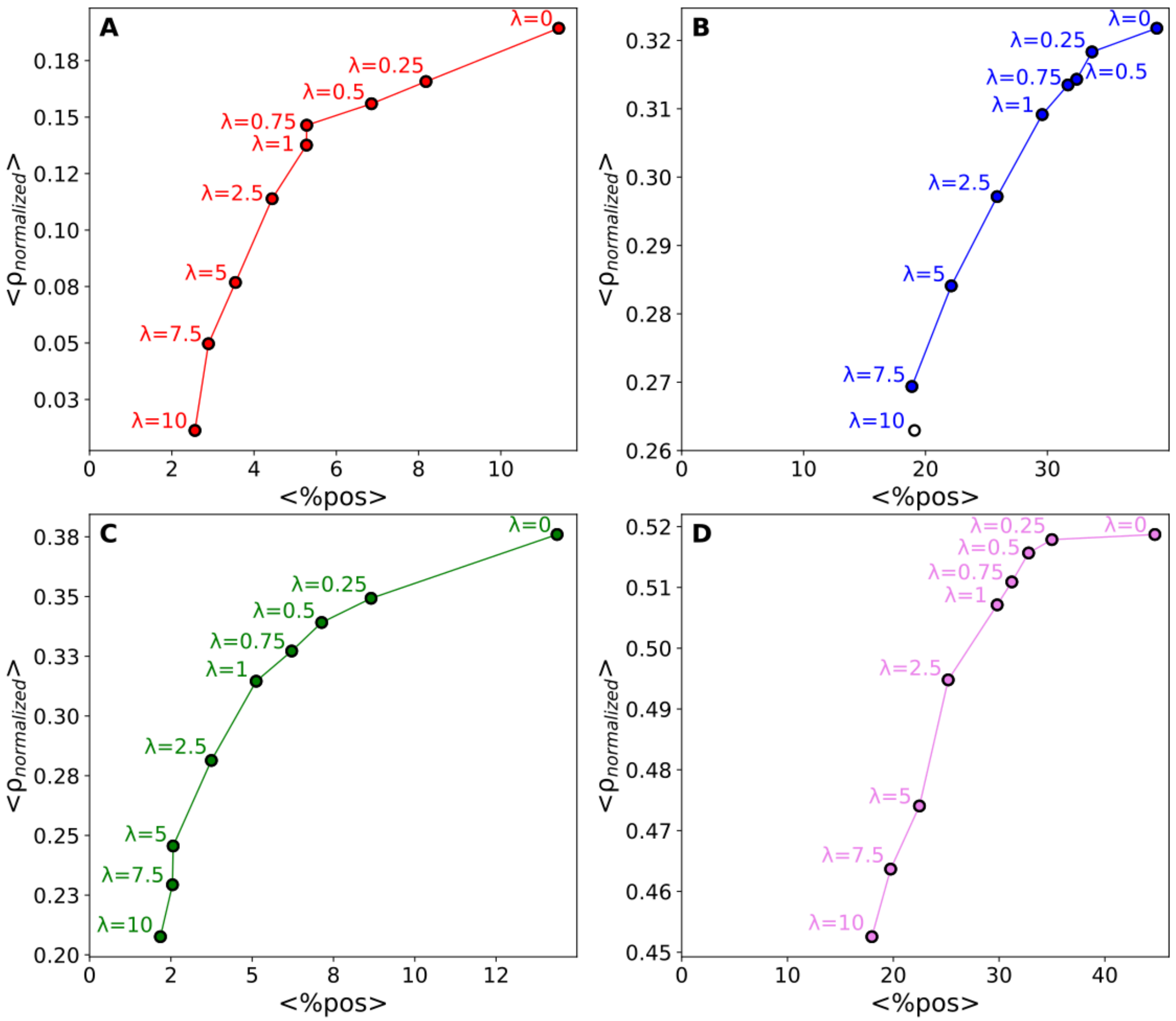


**Figure S1.** Multi-objective optimization between average fraction of Spearman correlation coefficient (ρ) improvement and percentage of alignment positions selected for each Monte Carlo (MC) positions selection algorithm and for a range of Lasso regularization coefficients (λ). For each protein family and each tested λ we ran the position-selection pipeline and recorded (i) the percentage of alignment positions retained by the method (x-axis) and (ii) The average fraction of improvement in ρ relative to the single site (SS) method as $<\rho_{normalized}> = <((\rho_i-\rho_{SS})/(1-\rho_{SS}))>$ (y-axis). Panels correspond to the algorithmic variants evaluated. (**A**) MC-SS, (**B**) MC-E, (**C**) AC-MC-SS, (**D**) AC-MC-E. This figure summarizes results across the 13 protein families analyzed.

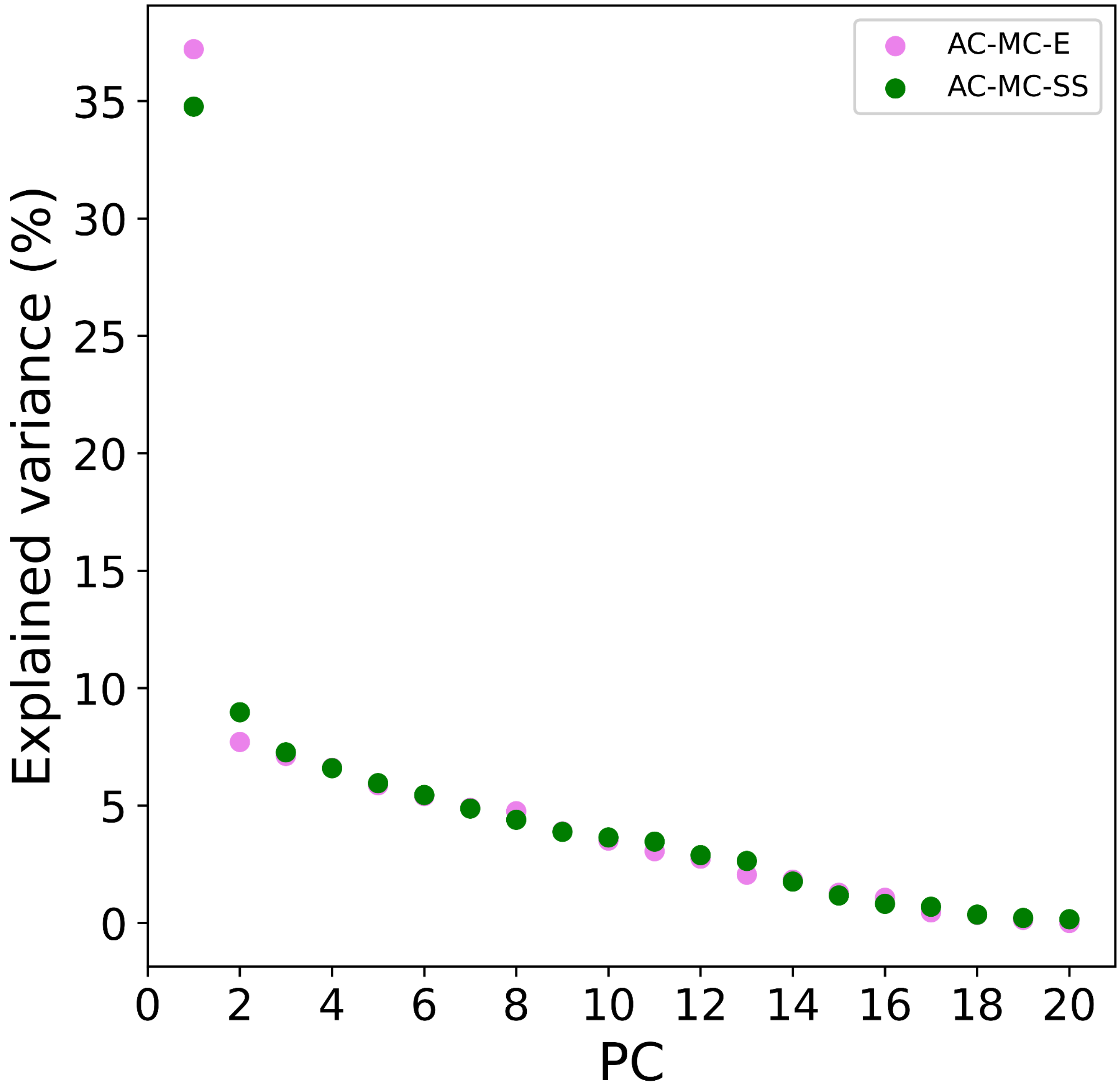


**Figure S2. Selection of optimal number of PCs by a Scree plot.** Explained variance calculated for each principal component for the selected positions of the AC-MC-SS (green) and AC-MC-E methods (violet). The optimal number of PCs was selected by keeping the ones to the left of the elbow of the curve, where the slope distinctly changes.

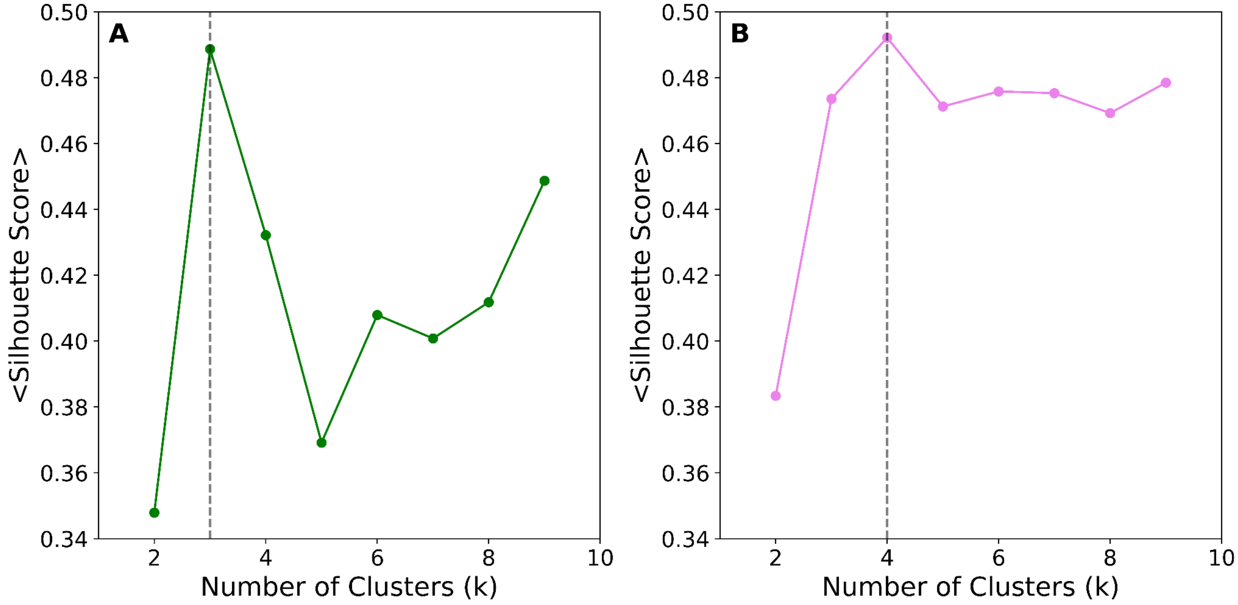


**Figure S3.** K-means clustering quality evaluation via silhouette analysis. Average silhouette scores calculated across a range of cluster numbers for the AC-MC-SS (**A**) and AC-MC-E (**B**) methods. The optimal number of clusters (k, indicated by the dashed vertical line) was selected by maximizing the global average silhouette score, which quantifies sample cohesion within assigned clusters relative to its separation from neighboring clusters.

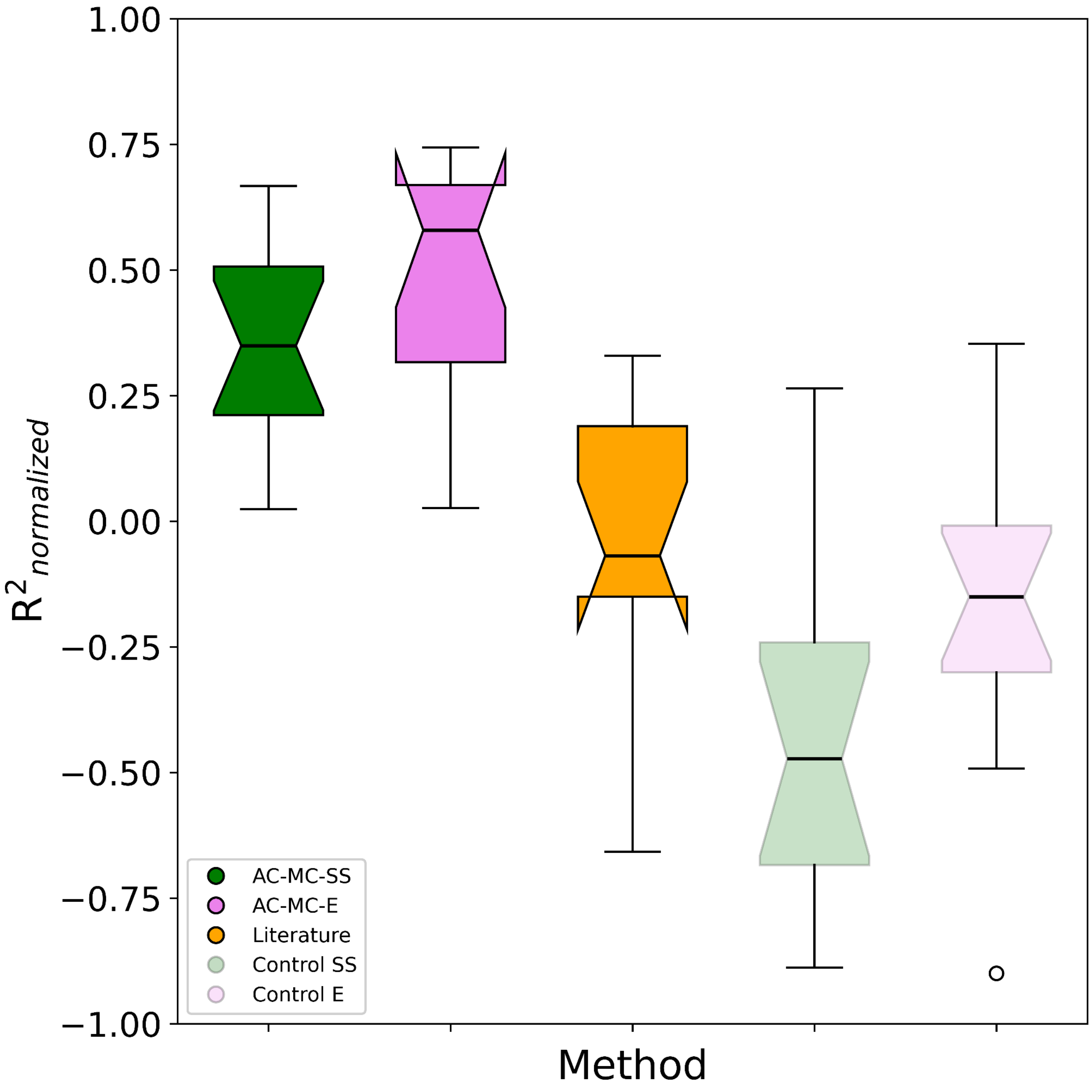


**Figure S5.** Box-plot summary of the distribution of average fraction of improvement in $R^2$ across case studies for AC-MC-E (violet), AC-MC-SS (green), their respective controls, and Literature (orange). Notches show the 95% confidence interval of the median: if two notches do not overlap, their medians differ significantly at approximately $\alpha = 0.05$. The y-axis shows the average fraction of improvement in $R^2$ relative to the single site (SS) method as $\langle R^2_{normalized} \rangle = \langle ((R^2_i - R^2_{SS})/(1-R^2_{SS})) \rangle$. Controls were generated by using an equivalent number of randomly selected positions or position pairs.

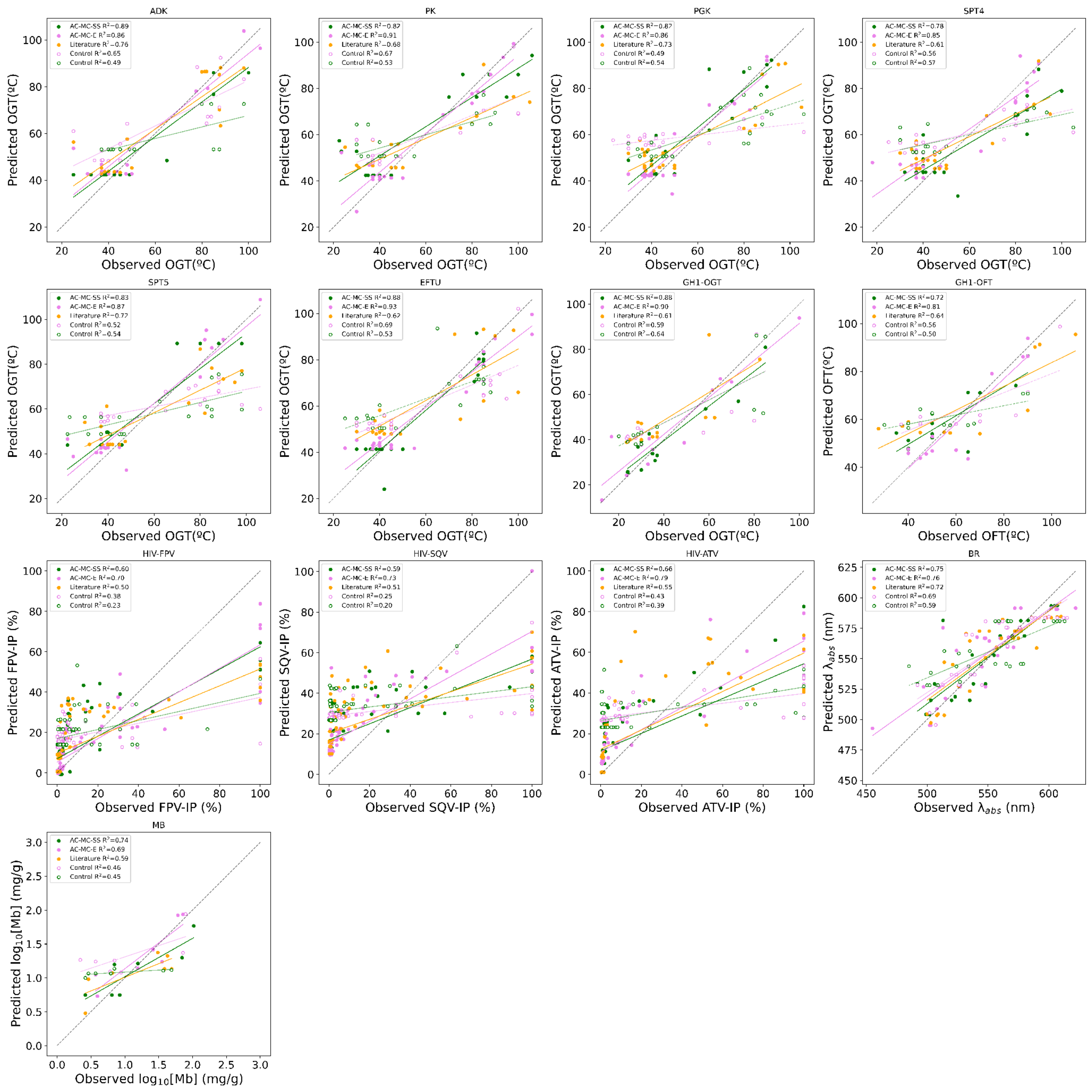

**Figure S6. Predicted vs. observed phenotypes across case studies**. Predictions based on positions selected by AC-MC-SS (green), AC-MC-E (violet), and literature data (orange). Solid lines represent linear fits for each method variant, while dashed green and violet lines (with unfilled markers) represent linear fits for their respective control calculations. Controls were generated by using an equivalent number of randomly selected positions or position pairs to calculate predictions. The dashed black line indicates the identity line (y=x).

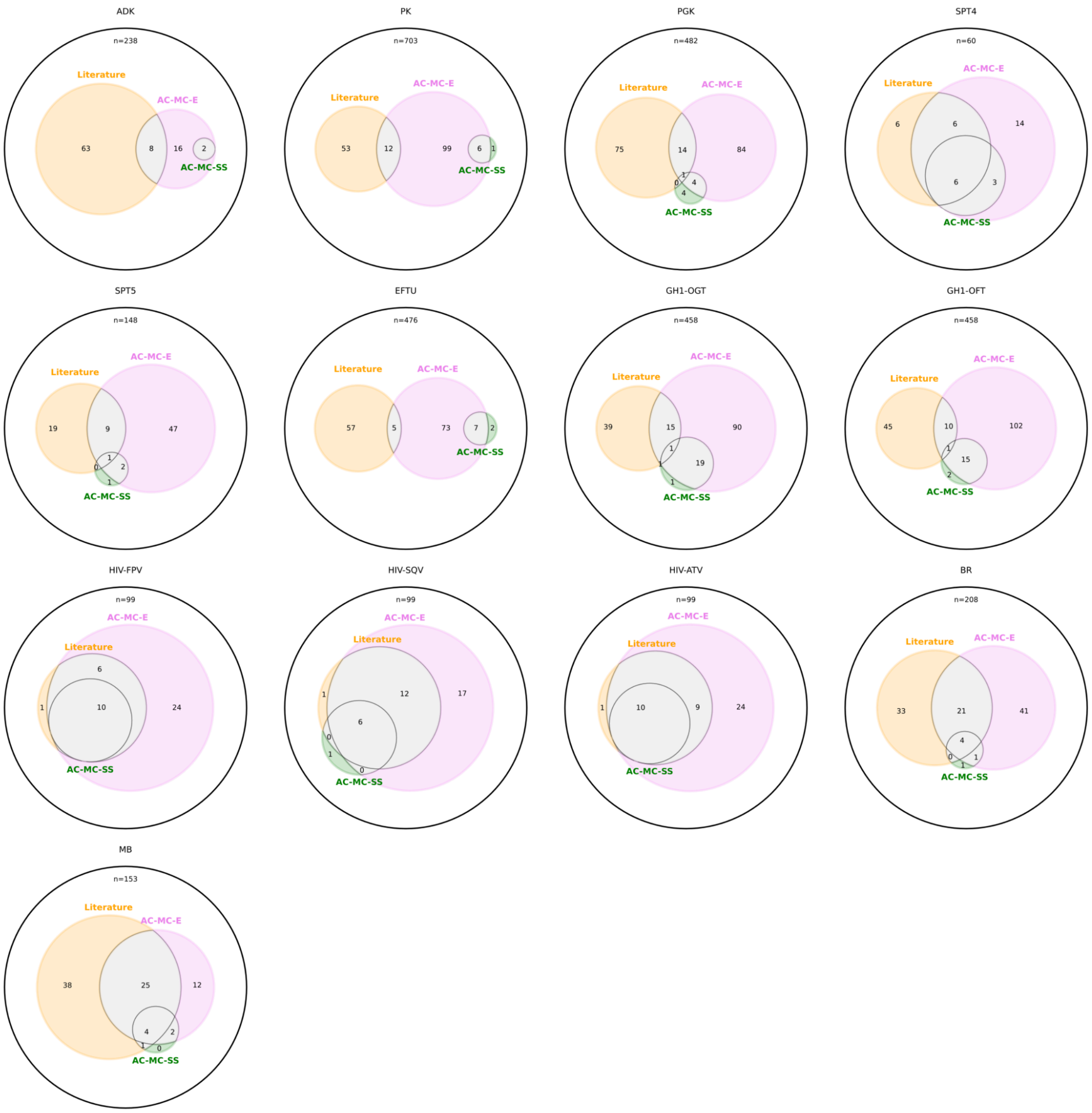


**Figure S7. Venn diagrams showing the overlap among alignment positions selected by each method for each multiple sequence alignment (MSA).** The black outer circle denotes the effective alignment universe (total number of alignment columns after excluding gap-only columns). Colored circles indicate the sets of selected positions: AC-MC-E (violet), AC-MC-SS (green), and literature positions (orange) . Numbers within each region report the size of each unique and shared subset of selected positions for that MSA.

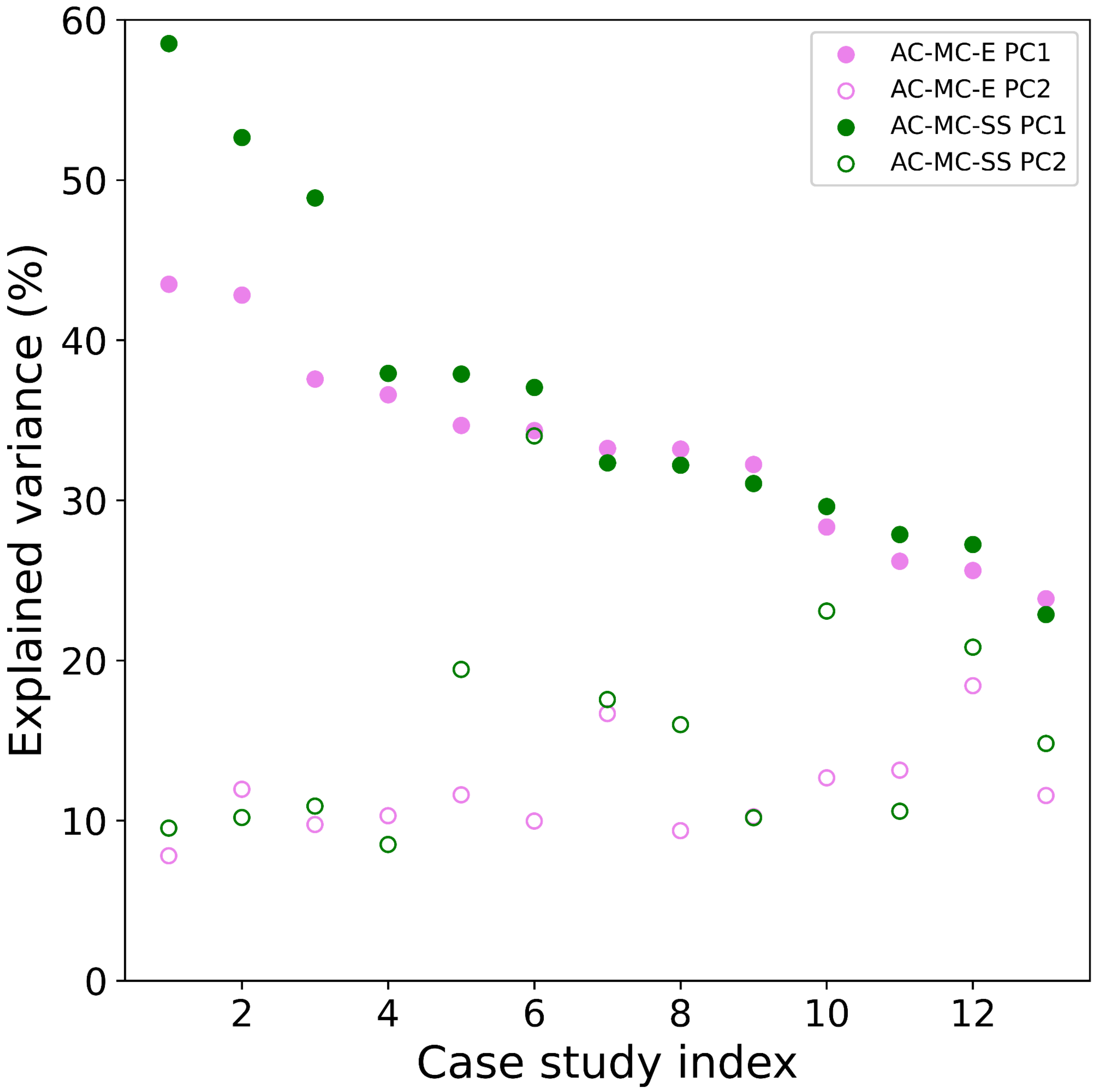


**Figure S8. Per case PCA explained variance.** Explained variance of the first two principal components (PC1, PC2) for each case study for the selected positions of AC-MS-SS (green) and AC-MS-E (violet) methods. Y-axis is sorted following the PC1 values in decreasing order for each method.

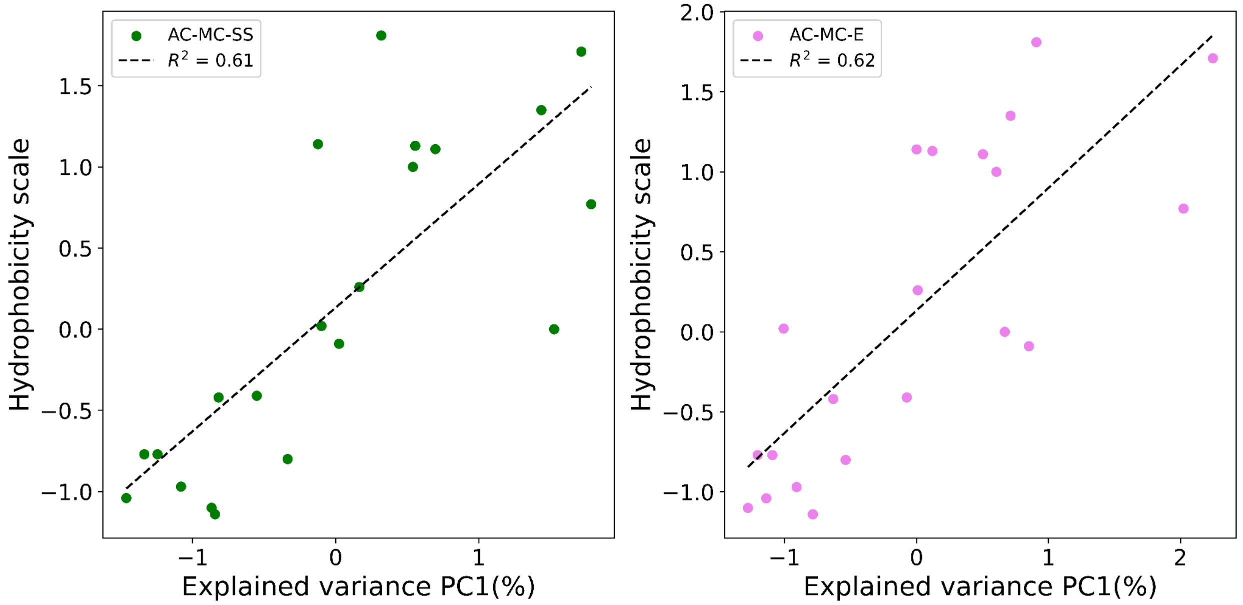


**Figure S9.** Correlation between PC1 explained variance and Hydrophobicity scale from Cid et al. 1992.
